# Ecological factors alter how spatial overlap predicts viral infection dynamics in wild rodent populations

**DOI:** 10.1101/2024.06.19.599763

**Authors:** Janine Mistrick, Jasmine S.M. Veitch, Katherine E. Wearing, Shannon M. Kitchen, Samuel Clague, Stephanie Du, Matthew Michalska-Smith, Brent C. Newman, Tarja Sironen, Clayton E. Cressler, Richard J. Hall, Sarah A. Budischak, Kristian M. Forbes, Meggan E. Craft

## Abstract

Spatial overlap between animals in wildlife populations can have important implications for pathogen transmission. Ecological factors and animal demographic traits can influence animal space use and spatial overlap, but it is unclear how these interactions drive pathogen transmission. We experimentally manipulated wild bank vole populations via resource supplementation and anthelmintic treatment. Using network analysis, we investigated the relationship between spatial overlap and infection likelihood of an endemic zoonotic hantavirus, including how vole sex and reproductive status interact with spatial behaviour to affect infection likelihood. Spatial overlap in a previous month drove the likelihood of current hantavirus infection, and food supplementation and anthelmintic treatment altered the effects of spatial overlap on infection likelihood. Vole sex and reproductive status were important factors determining whether spatial overlap increased or decreased the likelihood of hantavirus infection and interacted with resource supplementation and anthelmintic treatment, generating different infection dynamics in each treatment. Our research provides rare empirical evidence linking previous spatial overlap to current infection status in wildlife populations, with implications for understanding disease dynamics and persistence as well as developing effective management efforts. We further highlight the importance of incorporating variation in ecological factors and host demography when studying pathogen transmission in wildlife systems.

## INTRODUCTION

Animal behaviour, including space use and spatial overlap with individuals of the same and different species, is an important driver of transmission patterns of wildlife diseases. However, variation in ecological factors such as food resources and coinfection can influence pathogen transmission through both host space use and contact behaviour and within-host factors (1–3). Given that spatial overlap of wildlife, domestic animals, and humans is a key component of pathogen spillover (4), understanding the ecological drivers of transmission patterns in wildlife has crucial implications for public health (5) and conservation (6,7). To advance our understanding of transmission in wildlife populations, field experiments are necessary to test the effects of realistic natural variation in factors such as food resources and coinfection on the links between animal spatial behaviour and pathogen transmission (8).

Resource availability (1,9) and seasonal variation (10) can impact pathogen transmission by influencing animal space use and spatial overlap behaviour (3). For instance, low or patchy local food availability may increase host space use due to increased foraging behaviour, facilitating pathogen spread over greater distances (11). Increased spatial overlap in areas of high food availability can create opportunities for transmission and increase the prevalence of directly and indirectly transmitted pathogens (12,13). Moreover, spatial overlap among conspecifics and between species can also change seasonally due to resource availability (14) or social behaviours (e.g., mating, migration, co-denning 15,16) which can drive seasonal transmission patterns in wildlife populations (17).

Coinfection with macroparasites can impact microparasite transmission in wildlife via within-host effects on immune responses (2) or by altering animal space use and spatial overlap (18). Although parasites have well-established effects on animal behaviour (19), the effects of infection on animal space use and spatial overlap, and thus pathogen transmission potential, are poorly understood in wildlife. Infected animals may demonstrate sickness behaviours such as self-isolation (20), and costs of infection could decrease energetic activities such as movement (21,22) and mating, that could decrease space use and overlap (23). Uninfected animals may also avoid infected conspecifics or pathogen-contaminated landscapes, decreasing spatial overlap (24,25).

Ecological factors can also interact with animal demographic traits (e.g., sex, reproductive status) to influence space use and spatial overlap, and thus transmission opportunities within populations. Food availability and seasonality can influence demography at the population level by driving birth pulses (26,27) and increasing population density. Birth pulses can increase pathogen transmission by providing an influx of susceptible individuals and increasing pathogen exposure via greater spatial overlap. Population density can also alter individual-level demographic traits, such as the timing of sexual maturation (28). Reproductive animals often have larger space use (29) or more numerous interactions (30) compared to their non-reproductive conspecifics, providing more opportunities for transmission.

Despite the potential for ecological and demographic factors to influence animal space use and overlap, we lack empirical evidence of how these factors interact to influence individual infection risk, and thus population-level transmission (31). Social network analysis has become increasingly popular in the study of wildlife disease dynamics (32–34) to quantify how network structure (35,36), animal social behaviours (37–39), and demographic traits (36,40) influence population-level transmission dynamics. Many studies have used networks of contacts or overlap in wildlife to simulate or infer potential effects on transmission dynamics (reviewed in 31,41); however, empirical evidence in wildlife linking network connections with infection is much more limited (though a large body of work has been conducted in wild lizards 42–46). Moreover, such studies rarely incorporate variation in ecological factors such as food resources or coinfection. Integrating these ecological drivers into network analyses could enhance our understanding of drivers of seasonal and spatial variation in infection risk, with potential implications for disease management such as targeted vaccinations (17).

Controlled field experiments provide vital opportunities to quantify how ecological factors influence spatial interactions of animals and their consequences for pathogen transmission (41,47). Rodents and their endemic pathogens provide a tractable model system for field experiments investigating the effects of ecological factors such as food resources and macroparasite infection on microparasite transmission (13,48,49). For this study, we leveraged a well-characterised rodent-virus system: bank voles (*Clethrionomys glareolus*, previously *Myodes glareolus*) and Puumala hantavirus (PUUV). Bank voles are the reservoir host of the zoonotic microparasite PUUV, which is endemic in their populations, and are also commonly infected with macroparasites (helminths such as *Heligomosomoides glareoli*, 50). Bank vole populations, particularly in Fennoscandia, exhibit seasonal and multiannual cycles of population density and associated fluctuations in PUUV prevalence, with peak prevalence up to approximately 50% (51,52). PUUV is shed by voles in urine, faeces, and saliva, and transmission to conspecifics and other species occurs predominantly through environmental exposure; infectious virus can persist outside of the host for up to two weeks under laboratory conditions and potentially longer in some natural settings (53,54). Further, bank vole space use behaviour is well-documented, including how space use varies by sex, reproductive status, and season (29,55–57).

We experimentally manipulated food resources and helminth infections in wild bank vole populations and longitudinally measured vole spatial overlap and hantavirus infection to address the following research questions: 1) Does spatial overlap drive the likelihood of Puumala hantavirus infection in bank vole populations? 2) Is the relationship between spatial overlap and hantavirus infection likelihood influenced by food availability or helminth coinfection? and 3) How do vole sex and reproductive status interact with space use and spatial overlap to affect likelihood of hantavirus infection? Because infectious PUUV persists for around two weeks in the environment and seroconversion of infected voles can take days to weeks, we hypothesised that spatial overlap of a vole with conspecifics in a previous month would predict current infection likelihood. We also expected that vole space use would be influenced by food supplementation and anthelmintic treatment (58) and that demographic traits would be important drivers of the relationship between spatial overlap and likelihood of infection. We predicted that increases in spatial overlap would more strongly increase infection likelihood in demographic groups that are more tolerant of overlap (i.e., non-reproductive voles, reproductive males) compared to those that are less tolerant of overlap (i.e., reproductive females). We further hypothesised that changes in space use due to food supplementation or anthelmintic treatment would ultimately lead to different relationships between spatial overlap and hantavirus infection likelihood in each treatment.

## METHODS

### Study location and experimental design

A replicated two-factor field experiment was conducted in the boreal forests of southern Finland (61.0775°N, 25.0110°E) where bank voles are the dominant rodent species. The study design is summarised below and has been previously described in greater detail (59). Twelve study sites were established in old-growth spruce forest at least 2 km apart to prevent vole dispersal between sites. At each site, 61 traps were arranged in a standardised trapping grid (100 m x 100 m; with 10 m between grid rows and columns) to administer experimental treatments and monitor the vole population therein (**Figure 1A**). Sites were assigned one of four treatment pairings: both food supplementation and anthelmintic treatment (“fed-deworm”; “F-D”); food supplementation only (“fed-control”; “F-C”); anthelmintic treatment only (“unfed-deworm”; “U-D”); or no manipulation (“unfed-control”; “U-C”) with each treatment replicated at three sites. Food supplementation sites received a feed mix of rodent pellets and sunflower seeds roughly evenly distributed over the trapping grid every two weeks from May through November. At deworm sites, voles received an oral dose of 10 mg/kg Ivermectin and 100 mg/kg Pyrantel (effective at treating infection with larval and adult helminths, 60) at their first capture each month. At control sites, voles received a matching weight-based dose of sugar water (17.5% sucrose solution) at their first capture each month.

**Figure 1.**
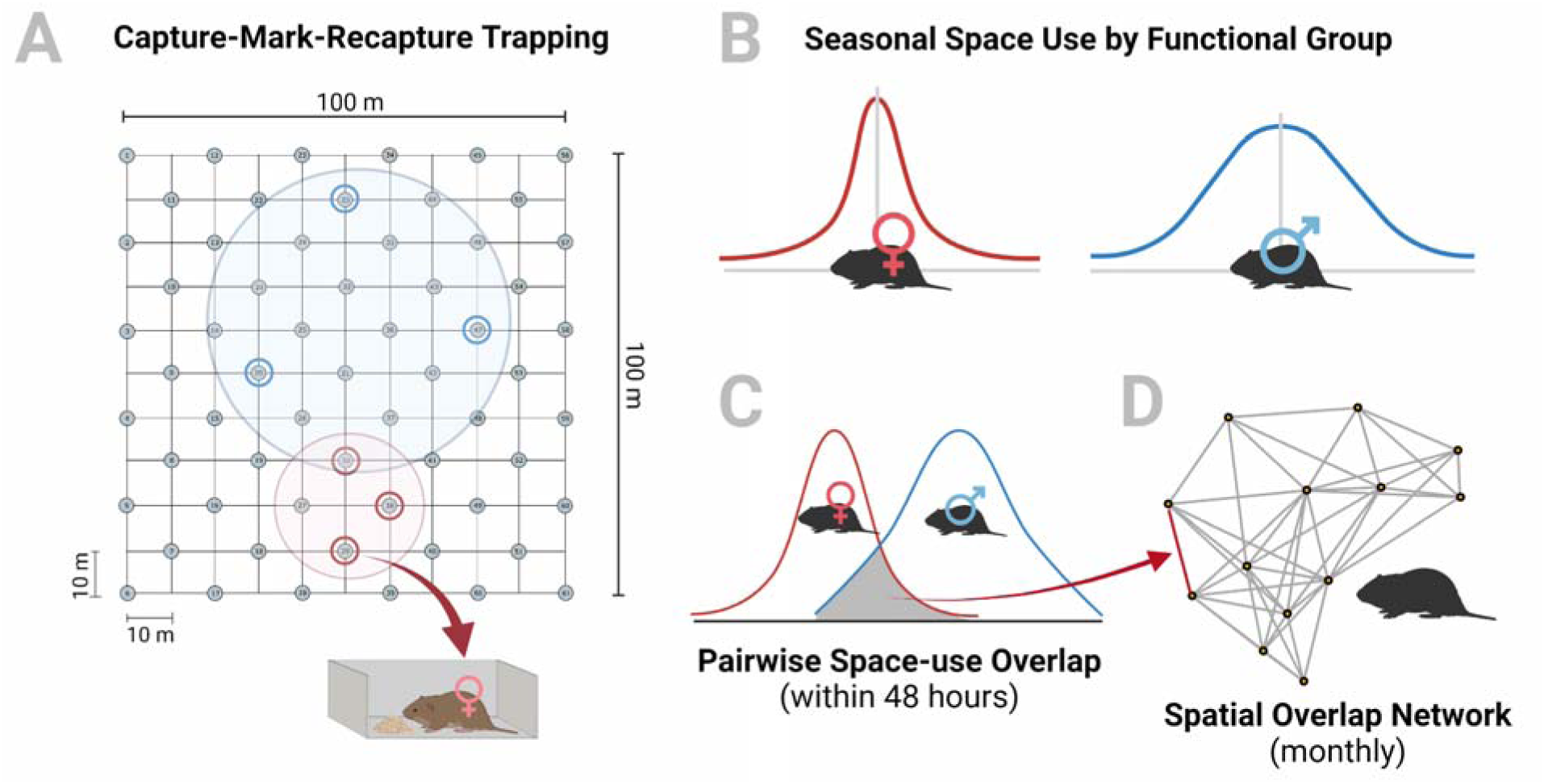
Conceptual figure of data collection and analysis. A) Voles were monitored via capture-mark-recapture methods and the trapped locations of individuals were recorded. B) Trapped locations were pooled across individuals by functional group (combination of sex and reproductive status), treatment, and season (summer or autumn). Space use was characterised for each group as the probability of capturing a vole of that group with increasing distance from the centroid of their trapped locations. This probability was assumed to be equal in all directions from the centroid, generating a circular kernel of space use. C) Space-use overlap between pairs of voles was estimated each month based on the overlap of the kernels for each vole’s functional group drawn around the centroids of their respective trapping locations. D) The amount of pairwise space-use overlap between voles was used to inform the edge weights of spatial overlap networks for voles at each site in each month June-October.

### Rodent monitoring

Vole populations were longitudinally monitored via capture-mark-recapture methods over two consecutive years (2021 and 2022). At each site, voles were trapped and sampled every four weeks from May-October (i.e., once per month for six months). Each month, traps were set for 48 hours and checked approximately every 12 hours (four checks total). Upon first capture, all voles were injected with a Passive Integrated Transponder (PIT tag) for unique identification. At their first capture in a month, the trap ID, PIT tag number, sex, and reproductive status of each vole were recorded. Voles were categorised as reproductive (males: scrotal testes; females: perforate vagina, visible nipples, or pregnant) or non-reproductive. Once per month, a faecal sample and a blood sample were collected from each vole. A faecal sample was collected from each vole to assess the efficacy of the oral anthelmintic treatment. A blood sample (up to 150 μL) was collected from the retro-orbital sinus of all captured voles >10 g using heparinized capillary tubes (Hirschmann, Germany). Whole blood samples were maintained in a cooler on ice until they were centrifuged and the serum separated. Serum was frozen at –20 °C until samples were screened for hantavirus infection (see below). If a vole was recaptured during the 48 hours of trapping in a month, only the trap ID and PIT tag number were recorded at subsequent captures and no samples were collected.

### Pathogen diagnostics

Vole hantavirus infection status was determined by detecting PUUV-specific IgG antibodies in blood serum via immunofluorescence assays (IFA), as previously described (61). In brief, serum samples were diluted in phosphate-buffered saline and incubated on slides treated with PUUV-infected cells, after which the slides went through a series of washes to remove unbound sample. Fluorescent polyclonal rabbit anti-mouse FITC conjugate was then incubated on the slides and slides were washed again to remove unbound conjugate. Slides then were viewed using a fluorescence microscope to detect reactive antibodies. Hantavirus infections in bank voles are chronic, thus the presence of IgG antibodies is used to indicate current infection (53,62).

A small subset of voles (21 individuals [totalling 50 unique captures]) were found to have inconsistent IFA results between subsequent samples (e.g., positive test followed by one or more negative tests, alternating negative and positive tests) and were removed from the dataset and excluded from downstream analysis. The majority of these (19/21 voles) were young animals (mean body mass 12.7g ± 1.6 standard deviation [range 10.5-16g]) that were positive at first capture and negative at subsequent capture(s). Maternal antibodies can persist in young bank voles for up to 80 days (63) and are indistinguishable from true infection via IFA, which is a likely source of inconsistent IFA results in individuals that were first captured at a young age.

Helminth infection intensity (measured as eggs per gram of faeces [EPG]) was quantified for each vole at each capture using a salt flotation method (64). This was used to quantify helminth infection status and infection intensity within the vole populations and confirm the efficacy of the anthelmintic treatment. The efficacy of the anthelmintic treatment was analysed separately in each year of the study. Generalised linear mixed-effects models (binomial family, logit link; lme4 R package, 65) were used to model helminth infection status in all sampled voles. Linear mixed-effects models (lmerTest R package, 66) were used to model helminth infection intensity (natural log-transformed EPG) in all infected voles. Both models were parameterised with treatment group (control/deworm), treatment stage (pre-treatment [the first capture of a vole] / post-treatment [all subsequent captures]), and their interaction as main effects and vole ID as a random effect.

### Vole space use & spatial overlap networks

The capture-mark-recapture data were used to characterise vole space use by season and quantify spatial overlap (defined here as mutual use of habitat, either concurrently or sequentially) within a month at each site. Vole captures in May, after the winter period, were very low (0-5 voles per site) so May data were excluded from analyses. Space use of reproductive bank voles changes between the summer breeding season (June-August) and autumn non-breeding season (September, October) in southern Finland (29,67) and previous research has established that vole space use varies by sex, reproductive status, and treatment between summer and autumn (58). Space use was therefore characterised by season by aggregating capture data across months for the summer and autumn. Space-use overlap between pairs of voles was measured in a shorter time frame (within the 48-hours of trapping in a month) to approximate opportunities for environmental transmission of PUUV. Spatial overlap networks were constructed to visualise and quantify the pairwise space-use overlap between voles at a site; one network was constructed for each of the five months when trapping was conducted (June-October).

Demographic traits such as sex and reproductive status can be used to categorise a population into functional groups, defined here to mean: population subgroups that are similar in their behaviour, physiology, and immunology (68–70). Conducting epidemiological analysis at the level of functional groups provides the biological context to draw more realistic conclusions about differences in contacts and pathogen prevalence within the population (71). Reproductive status was summarised across a season, such that a vole that was “reproductive” in any month June-August was classified as reproductive in summer. In autumn, voles were classified as reproductive if reproductive anatomy was observed in September or October or if the vole was classified as reproductive in the summer. Within a season, voles were then classified into four functional groups: reproductive males, reproductive females, non-reproductive males, and non-reproductive females.

The methods used herein to characterise seasonal space use and estimate monthly spatial overlap were developed by Wanelik & Farine (72) and have been proposed as a way to detect biological effects of spatial overlap even from sparse trapping data where observations per individual may be limited or heterogeneous. Detailed methods have been previously described (59) but are outlined here as follows. In order to calculate the seasonal space use of each individual vole, many of whom were only captured once or repeatedly in the same trap, voles were grouped and the captures of all individuals in the group were used to characterise an average seasonal space use kernel for a vole of that group. Voles were grouped by year, season, treatment, and functional group. Seasonal space use was characterised separately for each of these groups based on a negative sigmoidal curve modelling the relationship between the capture probability for an average vole in that group and the increasing distance from the vole’s “seasonal centroid” (i.e., the weighted average trapped location, weighted by the frequency of capture in each trap, for a given vole in that season). The relationship between capture probability and distance was assumed to be the same in all directions from the centroid, generating a circular kernel of seasonal space use for each group (**Figure 1B**).

To estimate monthly spatial overlap for individual voles, the weighted average trapped location for a vole in a given month was used as its “monthly centroid” and the seasonal space use kernel for the season, treatment, and vole’s functional group was centred at that point. This was repeated for all voles observed at a site in a given month. In this way, a vole captured in July and August would have the same “summer” space use kernel in each month, but their space use could be centred at different locations, based on where on the trapping grid that vole was captured in July versus August. Space-use overlap was estimated between all pairs of voles each month based on the amount of overlap in their space use kernels (“pairwise space-use overlap”, **Figure 1C**). Values of pairwise space-use overlap between two voles could range from 0 (no overlap) to 1 (complete overlap). These values were used to define the edge weights in undirected, weighted spatial overlap networks (“igraph” R package, 73) where voles were represented as nodes (**Figure 1D**). One network was constructed per month (June-October) at each site in both years of the study. Only voles captured in a given month were included in the network. All values of pairwise space-use overlap were used to define network edges with no thresholding, which is generally preferable to picking an arbitrary weight threshold and removing edges of lower weight or using unweighted edges (74).

From these monthly spatial overlap networks, measures of weighted degree were used to quantify the amount of spatial overlap between a focal vole and its neighbours. Weighted degree (“individual spatial overlap”) is the sum of the all pairwise space-use overlap between a focal vole and each of its neighbours. Weighted degree was partitioned by sex (male/female degree), reproductive status (reproductive/non-reproductive degree), and functional group (reproductive male/reproductive female/non-reproductive male/non-reproductive female degree) to quantify a focal vole’s overlap with each population subgroup. These weighted degree measures were calculated for every vole observed at a site in a given month.

In addition to spatial overlap, the tendency for voles to visit different traps was quantified as a measure of exploratory behaviour which could influence transmission opportunities in ways that are not captured in spatial overlap (for more details, see **Supplement: Exploratory behaviour**).

### Models of infection likelihood by treatment

A series of models were used to investigate how spatial overlap between bank voles affected Puumala hantavirus infection likelihood. The timing of seroconversion of PUUV-infected bank voles is variable and often takes several weeks or more (52,75). Thus, because trapping was conducted over 48 hours and repeated every four weeks, infection that was acquired during one month would not be detected until the following month. Therefore, models were parameterized to investigate the effect of a vole’s spatial overlap in the previous month on its infection status in the current month. Using infection likelihood as a proxy to infer transmission, we assumed that exposure is the major driver of transmission and factors such as individual differences in susceptibility play a limited role. It was rare to capture a vole in two months that were not consecutive (e.g., June and August, but not July) but in the cases where that did occur (25/713 [3.5%] of recaptures), the previous capture was used to inform current infection. Previous capture was restricted to within a sampling year, such that July was the first month when infection status (as informed by June network position) was considered.

Mixed-effects logistic regression was used to model the relationship between a vole’s current hantavirus infection status (infected/uninfected) and their previous weighted degree (“lme4” R package, 65). A series of candidate models were fit to data from each treatment separately. Infection status was the response variable in all models. Each degree measure (weighted degree, weighted degree partitioned by sex, weighted degree by reproductive status, weighted degree by functional group) was used as the main effect(s) in a separate model. Degree measures of all voles, even those with very little overlap, were included in models without thresholding.

For each degree measure, the interaction of sex, reproductive status, and functional group with previous degree were explored, each in separate models, resulting in 12 potential candidate models per treatment (See **Results: Models of infection likelihood** and **Table S1** for subset of candidate models successfully fitted per treatment). Additional main effect variables used in all models were vole sex, reproductive status, and exploratory behaviour, the previous month and previous network size (corresponding to the previous degree measure), and sampling year. Site ID was included as a random effect to account for inherent variation among populations.

Numeric variables were scaled and centred to improve model convergence. “Previous month” was coded as a numeric variable for the unfed-control, unfed-deworm, and fed-deworm treatments. However, “previous month” was coded as a categorical variable (levels ordered chronologically) for the fed-control treatment to address singularity issues in model fitting. All candidate models per treatment were assessed by Akaike Information Criterion (AIC) and the most parsimonious was selected. From the best-fit model for each treatment, significant interaction terms were used to identify correlation between previous spatial overlap and current infection status (to address research question 1). If different interaction terms were significant in different treatments or the effect size of a single term changed between treatments, that would indicate that the relationship between spatial overlap and infection likelihood was influenced by the manipulated ecological factor (research question 2). Effect sizes that differed within a treatment based on the sex or reproductive status of the focal and neighbour voles would indicate that demographic traits interacted with spatial overlap to affect infection likelihood (research question 3). All analyses were conducted in programme R version 4.3.2 (76).

## RESULTS

Across the two years of the study (May-October 2021 and 2022), we captured 1,518 unique voles, 929 [61%] of which were recaptured. Spatial overlap networks were constructed with all voles captured in June-October with recorded sex and reproductive status data: 1,129 captures (742 unique voles) in 2021 and 1,131 captures (744 unique voles) in 2022. Of these, 1,367 voles were tested for PUUV antibodies via IFA (n=683 in 2021; n=694 in 2022 – 10 animals were captured in both years). In 2021, 30.3% of tested voles were hantavirus positive. In 2022, 16.6% of tested voles were hantavirus positive. To model hantavirus infection likelihood, only animals with network and hantavirus data and captures in at least two separate months were used, resulting in datasets of 346 captures (237 unique voles) in 2021 and 367 captures (263 unique voles) in 2022.

Faecal egg count data was collected for 1,010 captures in 2021 and 1,007 captures in 2022. Of these, helminth eggs were detected in 434 samples (43%) and 262 samples (26%), respectively. In both years of the study, anthelmintic treatment decreased helminth infection prevalence at subsequent captures (i.e., post-treatment) in treated voles compared to voles given a sugar water control (2021: odds ratio=0.455, p=0.008; 2022: odds ratio=0.495, p=0.047). Anthelmintic treatment decreased infection intensity in infected voles in the first year of the study (2021: {3=-0.74, p=0.004) but no effect was detected in the second year (2022: {3= –0.04, p=0.915).

### Vole space use & spatial overlap networks

In the summer breeding season (June-August), seasonal space use was generally greater for males than females in both reproductive and non-reproductive voles (**Figure 2**; **Figure 3A**), and, overall, greater for reproductive voles than non-reproductive voles (**Figure 2**; **Figure 3A** e.g., August). In the autumn non-breeding season (September-October), space use was more similar across functional groups (**Figure 3A**), though reproductive vole space use was slightly greater than that of non-reproductive voles (**Figure 2**). Reproductive vole space use decreased between summer and autumn while non-reproductive vole space use increased. These trends were consistent across treatments and study years. By treatment, food supplementation decreased space use in both seasons in 2021, but no effect of food supplementation was detected in 2022 (**Table 1**). In both years, anthelmintic treatment had no effect in summer, but increased space use in autumn (**Table 1**).

**Figure 2.**
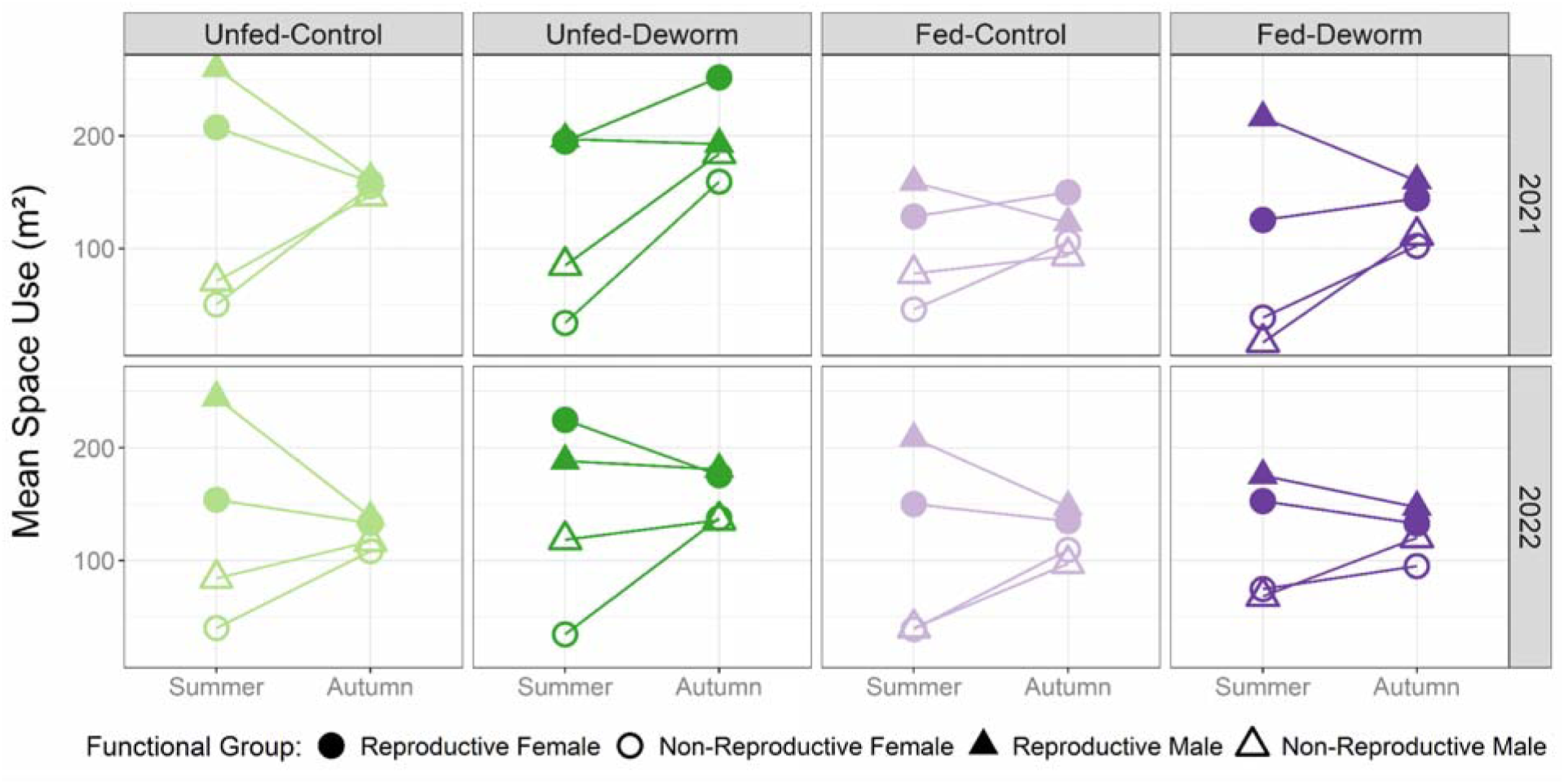
Vole space use by functional group in summer and autumn across treatments and study years. Reported values indicate area of space use where the probability of capture was >0.01 (measured in m^2^) based on the circular kernel of space use from the negative sigmoidal curve. Greater variation in space use across functional groups was observed in summer compared to autumn. Space use generally decreased for reproductive voles (filled shapes) and increased for non-reproductive voles (open shapes) between summer and autumn.

**Figure 3.**
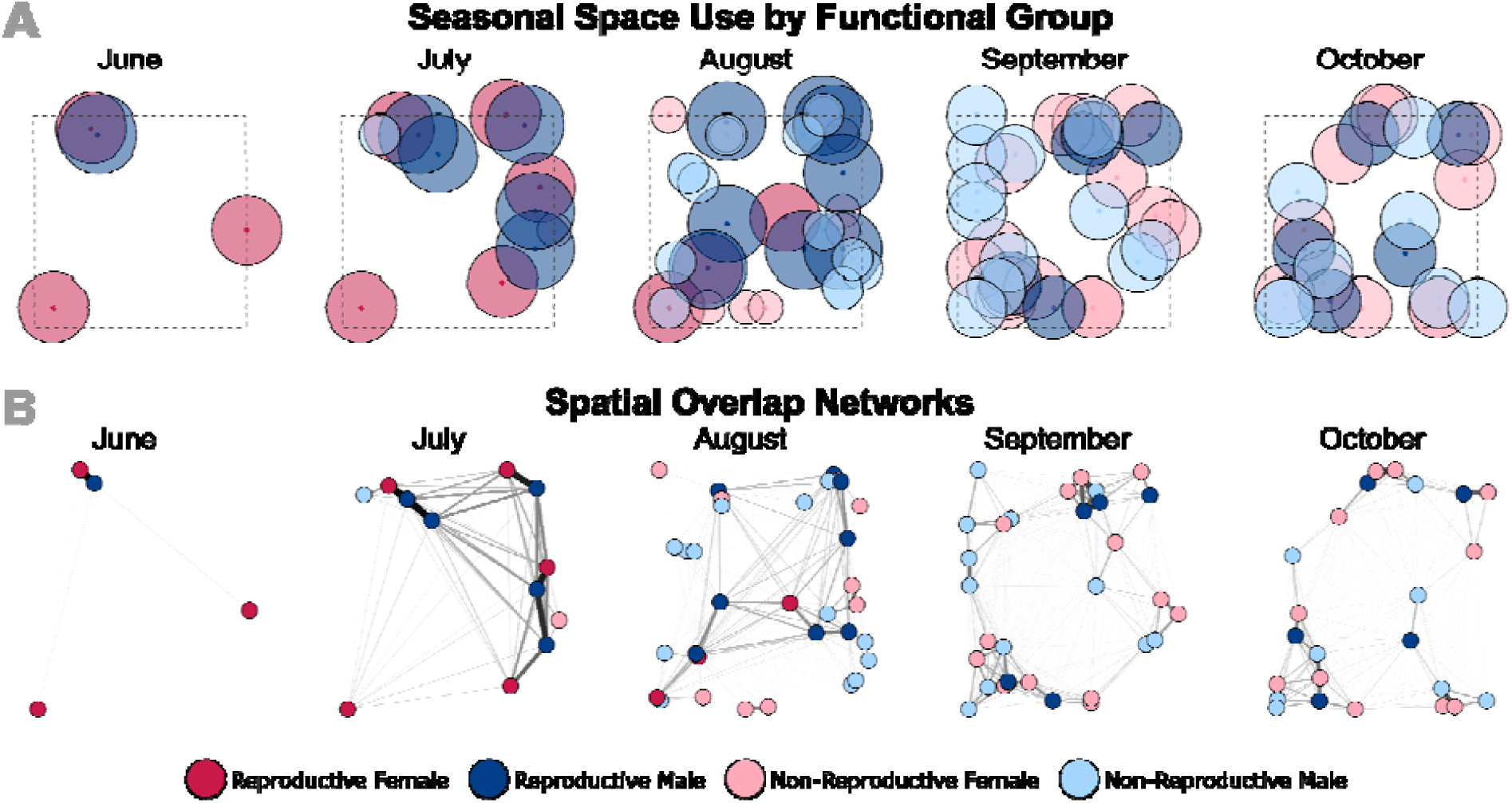
Seasonal space use and spatial overlap networks throughout the study period in 2021 for one of the investigated vole populations (unfed-control treatment, site “Helmipöllö”). A) Coloured circles indicate space use of voles captured at the site each month. Space use was characterised separately for each functional group across the summer breeding season (June-August) and across the autumn non-breeding season (September-October). Reproductive males (dark blue) had larger space use than reproductive females (red) in summer. Larger, darker circles in June-August indicate reproductive voles. Non-reproductive voles (smaller, lighter circles) first appear in July. Reproductive and non-reproductive vole space use was similar in autumn. B) Spatial overlap networks were constructed each month with edges weighted based on the amount of space-use overlap between pairs of voles (edge weight values scale 0 to 1). Network nodes are positioned in space to match the location of voles’ monthly centroids as seen in (A). Edges of higher weight (indicating greater overlap) are thicker and darker in colour.

**Table 1.** Parameter estimates describing the effects of food supplementation and helminth removal on bank vole space use. Four separate generalised linear models were constructed to predict how the probability of capturing a vole varied with the natural logarithm of distance from its seasonal centroid, indicating space use, in each year and season of the study. Food supplementation, helminth removal, vole sex, and reproductive status were all included as parameters influencing capture probability, but only the model estimates for the interaction of distance and the experimental manipulations (“food supp” and “deworm”) are shown. Bolded odds ratios reflect a significant effect of food supplementation or helminth removal on bank vole space use.

| Year | Season | Variable | Odds Ratio [95% CI] | p-value |
| --- | --- | --- | --- | --- |
| 2021 | Summer | ln(Distance) * **Food Supp** | **0.31 [0.16, 0.58]** | **p<0.001** |
|  |  | ln(Distance) * Deworm | 0.65 [0.33, 1.22] | p=0.18 |
|  | Autumn | ln(Distance) * **Food Supp** | **0.23 [0.09, 0.54]** | **p<0.001** |
|  |  | ln(Distance) * **Deworm** | **2.59 [1.22, 5.50]** | **p=0.013** |
| 2022 | Summer | ln(Distance) * Food Supp | 0.83 [0.48, 1.44] | p=0.51 |
|  |  | ln(Distance) * Deworm | 0.92 [0.49, 1.69] | p=0.78 |
|  | Autumn | ln(Distance) * Food Supp | 1.11 [0.44, 2.87] | p=0.83 |
|  |  | ln(Distance) * **Deworm** | **3.21 [1.27, 8.26]** | **p=0.014** |

Spatial overlap networks were constructed for June-October for all 12 sites in 2021 and 2022 with the exception of one site that had no animals in June 2022 so networks were only constructed for July-October. The observed spatial overlap networks generally showed high connectivity (many of the possible edges between individuals were realised; **Figure 3B**) but few of these edges were of high weight. Focusing on edges of higher weight, indicating a higher probability of overlap between pairs of voles, showed heterogeneous network structure with clusters of closely interacting voles, particularly in August-October. Distributions of weighted degree values each month were generally similar across treatments, though distributions tended to be more right skewed (indicating more voles with greater individual spatial overlap) in the fed treatments compared to unfed and in 2022 compared to 2021. Mean weighted degree was similar between treatments each month and was slightly higher in 2022 compared to 2021 (mean weighted degree ± standard deviation: 2021: 1.40 ± 0.88; 2022: 1.6 ± 1.05; **Figure S1**)

### Models of infection likelihood by treatment

The best-fit model to predict current hantavirus infection from previous spatial overlap for both the unfed-control and unfed-deworm treatments was the model with previous weighted degree partitioned by reproductive status and an interaction with sex. The best-fit model for both the fed-control and fed-deworm treatments was the model with previous weighted degree partitioned by functional group and interactions with both sex and reproductive status.

A low sample size of hantavirus-positive non-reproductive voles in three of the treatments (unfed-control, unfed-deworm, and fed-control) prevented us from fitting the full suite of candidate models to these datasets (for list of models fit by treatment see **Table S1**). In the unfed-control treatment, no hantavirus-positive non-reproductive voles were present in the dataset so all candidate models were fit without the “reproductive status” parameter. Overall, sample size was lower in the unfed-deworm treatment (n observations=110) due to low vole capture numbers as compared to the other three treatments (n observations U-C=183; F-C=189; F-D=231).

For the best-fit model for each treatment, we identified the significant interaction terms which indicated that previous spatial overlap was correlated with current infection status for voles of a given sex and reproductive status. In the unfed-control treatment, previous spatial overlap affected the likelihood of current hantavirus infection for male voles. Previous spatial overlap with non-reproductive voles decreased the likelihood of current infection in males while previous spatial overlap with reproductive voles increased the likelihood of infection (non-reproductive degree*male Odds Ratio (OR)=0.05, p=0.079, 95% CI [0.0, 1.41]; reproductive degree*male OR=1.74, p=0.025, 95% CI [1.07, 2.82]; **Figure 4A-B**; **Table S2**). In the unfed-deworm treatment, previous spatial overlap with non-reproductive voles increased the likelihood of current infection in female voles (non-reproductive degree*female OR=21.3, p=0.017, 95% CI [1.72, 266]; **Figure S2**; **Table S3**).

**Figure 4.**
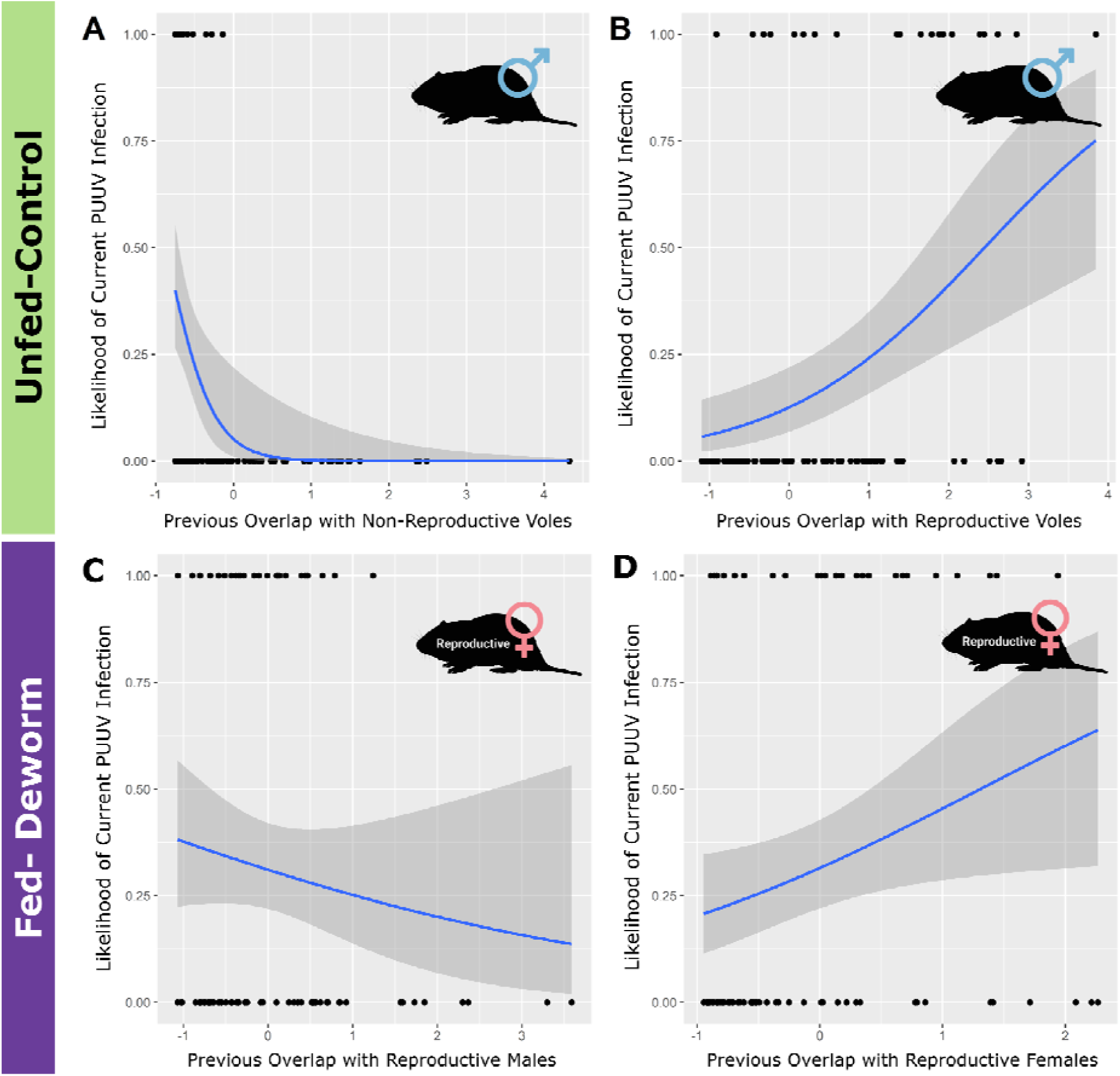
Correlation between the likelihood of current Puumala hantavirus (PUUV) infection and previous spatial overlap (where overlap values were scaled and centred). For male voles in the unfed-control treatment, previous overlap with (A) non-reproductive voles decreased infection likelihood and (B) reproductive voles increased infection likelihood. For reproductive female voles in the fed-deworm treatment, previous overlap with (C) reproductive male voles decreased infection likelihood and (D) reproductive female voles increased infection likelihood. Points indicate raw data.

In the fed-control treatment, previous spatial overlap with non-reproductive male voles increased the likelihood of current infection for reproductive females (non-reproductive male degree*reproductive*female OR=2.81, p=0.026, 95% CI [1.13, 6.97]; **Table S4**). We also found weak support for previous overlap with reproductive females decreasing the likelihood of current infection in non-reproductive voles (see **Supplement: Fed-Control Treatment; Figure S3**). In the fed-deworm treatment, previous spatial overlap affected the likelihood of current infection for reproductive females. Previous spatial overlap with reproductive males decreased the likelihood of current infection for reproductive females while previous spatial overlap with reproductive females increased the likelihood of current infection for reproductive females (reproductive male degree*reproductive*female OR=0.29, p=0.015, 95% CI [0.11, 0.79]; reproductive female degree*reproductive*female OR=4.22, p=0.005, 95% CI [1.54, 11.6]; **Figure 4C-D**; **Table S5**).

## DISCUSSION

We experimentally manipulated wild bank vole populations via food supplementation and anthelmintic treatment and investigated the effects of spatial overlap on infection likelihood of an endemic, environmentally transmitted hantavirus. Our findings demonstrate that spatial overlap with other voles drove hantavirus infection likelihood in subsequent months. Food supplementation and anthelmintic treatment altered the relationship between spatial overlap and infection likelihood, resulting in overlap correlating differently with hantavirus infection in each treatment. Further, the sex and reproductive status of overlapping voles influenced whether increased overlap increased or decreased infection likelihood. Broadly speaking, our results indicate that host demography and ecological context play a large role in determining how wildlife space use and spatial overlap influence infection risk.

Spatial overlap in the previous month drove the likelihood of current hantavirus infection, indicating a link between shared use of the environment and pathogen exposure. Most knowledge of how spatial overlap shapes environmental pathogen transmission has focused on key spatial locations where wildlife congregate, such as dens, nests, or refuges (35,44,77) or point sources of food or water (78,79). Studies exploring the role of spatial overlap more broadly have found mixed effects on transmission-related outcomes: spatial overlap predicted contact rates in racoons (80) and predicted tick loads and tick-borne blood parasite infection in a territorial lizard (45), but did not predict faecal-oral bacterial infection in other studies of lizards and giraffes (39,42). Our findings have contributed new evidence for how spatial overlap is related to infection likelihood of an environmentally transmitted pathogen in wild rodents. This adds support that spatial overlap could be beneficial in understanding transmission patterns in wildlife, though further studies across diverse species and pathogen transmission routes are needed to determine the generalizability of these patterns.

Across treatments, the relationship between spatial overlap and hantavirus infection was affected by food supplementation and anthelmintic treatment. We found that previous spatial overlap predicted current hantavirus infection likelihood for a different functional group under each experimental treatment. This complexity demonstrates that – even in a single host-pathogen system – ecological factors can generate variability in how spatial overlap influences infection likelihood. Previous research has similarly suggested that food availability (1,81) and helminth infection (2,82) can have highly variable effects on pathogen dynamics in wildlife. Moreover, interactions between food availability and helminth infection can interact with the host immune system to generate synergistic effects on pathogen dynamics (82,83). However, these studies largely focused on within-host aspects of transmission such as susceptibility and not on between-host aspects such as contacts and exposure. Few other studies have empirically tested if different ecological factors alter the associations between spatial overlap and infection likelihood, and thus our findings provide perspective which is often absent from studies of pathogen transmission.

Increased overlap between voles did not necessarily correlate with an increased likelihood of infection. Notably, the sex and reproductive status of focal voles and their overlapping neighbours interacted to determine whether increased spatial overlap increased or decreased infection likelihood. This heterogeneity points to an important, broader limitation of using spatial overlap to infer transmission opportunities: spatial overlap does not account for animal behaviour such as attraction or avoidance between individuals (47,74). While it is commonly assumed that the most connected individuals are at the highest risk of infection, social network studies of wildlife suggest that the type of interactions between individuals and their directionality may be more important to transmission than simply the total amount of exposure (37). Differences in social behaviour by sex or age class can also result in connections to certain conspecifics, e.g., males (45) or highly mobile juveniles (36), conferring a higher risk of infection than connections to other individuals. Leveraging data on social behaviour can thus provide insights into otherwise counterintuitive findings such as why increased overlap between male voles and reproductive voles increased infection likelihood, but increased overlap between male voles and non-reproductive voles decreased infection likelihood. Non-reproductive voles do not hold territories and instead “sit and wait” on the periphery of mature voles’ home ranges (Bujalska, 1988), resulting in limited opportunities for transmission. Moreover, maternal antibodies can protect young voles from infection for several months (Kallio, Poikonen, et al., 2006), making them an unlikely source of infection for conspecifics. On the other hand, reproductive animals have more interactions with conspecifics via mating and marking or defending territory and thus have had more opportunities to acquire and transmit pathogens. As such, when spatial overlap is used to approximate transmission, it is critical to consider the demographic traits of overlapping individuals, as this will determine the types of interactions associated with spatial overlap and whether they facilitate or limit pathogen transmission. In future research, simultaneous collection and analysis of both spatial and social data may help to illuminate the relative contributions of each and their effects on transmission dynamics (84).

Understanding how an animal’s spatial behaviour, sex, and reproductive status influence both their likelihood of infection and their role in spreading pathogens is central to understanding disease dynamics and persistence, as well as targeted disease management efforts such as vaccination and removal (34). However, if the type of individuals at highest risk of acquiring or transmitting infection varies based on the prevailing ecological conditions, this could decrease the efficacy of targeted control efforts in ecologically variable environments. Ultimately, our findings show that food resources and helminth infection can interact with animal sex and reproductive status to alter how spatial overlap affects infection likelihood of an environmental pathogen. This highlights the importance of incorporating biologically relevant variation in ecological factors and host demography into studies of infectious diseases in wildlife to more fully understand ecological drivers of pathogen transmission.

## ACKNOWLEDGEMENTS

We are grateful to our collaborators at the Lammi Biological Station and University of Helsinki: John Loehr, Janne Sundell, Esa-Pekka Tuominen, Joni Uusitalo, Tiina Tulonen, Matti Kotakorpi, Riitta Ilola, Jaakko Vainionpää, Tomas Strandin, and Sanna Mäki, and to the students who helped collect and process the field data: Alexis Beagle, Emilie Bonhomme, Muriel Chaudhri, Juliane Damaschke, Alyssa Dunn, Lucie Fornili, Mathilde Gaudillère, Ibrahim Koroma, Teemu Lemola, Finley Melnikoff, Eléonore Miston, Nathaniel Mull, Eunice Oh, Amy Schexnayder, Anni Simonen, Isabella Stark, and Léa Tambereau. Thanks to Brendan Haile, Sharon Jansa, and Susan Jones for providing feedback on this work. Figures were created with BioRender.com.

## USE OF ARTIFICIAL INTELLIGENCE

No AI technologies were used in the preparation of this manuscript.

## ETHICS

All live animal procedures were approved by the XXX Institutional Animal Care and Use Committee (IACUC #19105) and the Finnish Animal Ethics Board (ESAVI-17810-2019). Access to forest sites was provided by private landowners and by Metsähallitus Metsätalous Oy (MH 6302/2019).

## DATA, CODE, and MATERIALS

Vole capture and hantavirus infection data and the R code to run all analyses and generate all the figures will be made available through GitHub and the Dryad Digital Repository (85)

## REFERENCES

1. Becker DJ, Streicker DG, Altizer S. Linking anthropogenic resources to wildlife–pathogen dynamics: a review and meta-analysis. Ecol Lett. 2015;18(5):483–95.

2. Ezenwa VO. Helminth–microparasite co-infection in wildlife: lessons from ruminants, rodents and rabbits. Parasite Immunol. 2016;38(9):527–34.

3. VanderWaal KL, Ezenwa VO. Heterogeneity in pathogen transmission: mechanisms and methodology. Hawley D, editor. Funct Ecol. 2016 Oct;30(10):1606–22.

4. Plowright RK, Parrish CR, McCallum H, Hudson PJ, Ko AI, Graham AL, et al. Pathways to zoonotic spillover. Nat Rev Microbiol. 2017 Aug;15(8):502–10.

5. Plowright RK, Reaser JK, Locke H, Woodley SJ, Patz JA, Becker DJ, et al. Land use-induced spillover: a call to action to safeguard environmental, animal, and human health. Lancet Planet Health. 2021 Apr;5(4):e237–45.

6. Daszak P, Cunningham AA, Hyatt AD. Emerging Infectious Diseases of Wildlife--Threats to Biodiversity and Human Health. Science. 2000 Jan 21;287(5452):443–9.

7. Smith KF, Sax DF, Lafferty KD. Evidence for the Role of Infectious Disease in Species Extinction and Endangerment. Conserv Biol. 2006;20(5):1349–57.

8. Tompkins DM, Dunn AM, Smith MJ, Telfer S. Wildlife diseases: from individuals to ecosystems. J Anim Ecol. 2011;80(1):19–38.

9. Altizer S, Becker DJ, Epstein JH, Forbes KM, Gillespie TR, Hall RJ, et al. Food for contagion: synthesis and future directions for studying host–parasite responses to resource shifts in anthropogenic environments. Philos Trans R Soc B Biol Sci. 2018 May 5;373(1745):20170102.

10. Altizer S, Dobson A, Hosseini P, Hudson P, Pascual M, Rohani P. Seasonality and the dynamics of infectious diseases. Ecol Lett. 2006;9(4):467–84.

11. Becker DJ, Hall RJ, Forbes KM, Plowright RK, Altizer S. Anthropogenic resource subsidies and host–parasite dynamics in wildlife. Philos Trans R Soc B Biol Sci. 2018 May 5;373(1745):20170086.

12. Cross PC, Edwards WH, Scurlock BM, Maichak EJ, Rogerson JD. Effects of Management and Climate on Elk Brucellosis in the Greater Yellowstone Ecosystem. Ecol Appl. 2007;17(4):957–64.

13. Forbes KM, Henttonen H, Hirvelä-Koski V, Kipar A, Mappes T, Stuart P, et al. Food provisioning alters infection dynamics in populations of a wild rodent. Proc R Soc B Biol Sci. 2015 Oct 7;282(1816):20151939.

14. VanderWaal KL, Gilbertson MLJ, Okanga S, Allan BF, Craft ME. Seasonality and pathogen transmission in pastoral cattle contact networks. R Soc Open Sci. 2017 Dec 6;4(12):170808.

15. Hirsch BT, Reynolds JJH, Gehrt SD, Craft ME. Which mechanisms drive seasonal rabies outbreaks in raccoons? A test using dynamic social network models. J Appl Ecol. 2016;53(3):804–13.

16. Strona ALS, Levenhagem M, Leiner NO. Reproductive effort and seasonality associated with male-biased parasitism in *Gracilinanus agilis* (Didelphimorphia: Didelphidae) infected by *Eimeria* spp. (Apicomplexa: Eimeriidae) in the Brazilian cerrado. Parasitology. 2015 Jul;142(8):1086–94.

17. Rushmore J, Caillaud D, Hall RJ, Stumpf RM, Meyers LA, Altizer S. Network-based vaccination improves prospects for disease control in wild chimpanzees. J R Soc Interface. 2014 Aug 6;11(97):20140349.

18. Peacock SJ, Krkošek M, Lewis MA, Molnár PK. A unifying framework for the transient parasite dynamics of migratory hosts. Proc Natl Acad Sci. 2020 May 19;117(20):10897–903.

19. Ezenwa VO, Archie EA, Craft ME, Hawley DM, Martin LB, Moore J, et al. Host behaviour– parasite feedback: an essential link between animal behaviour and disease ecology. Proc R Soc B Biol Sci. 2016 Apr 13;283(1828):20153078.

20. Hawley DM, Altizer SM. Disease ecology meets ecological immunology: understanding the links between organismal immunity and infection dynamics in natural populations. Funct Ecol. 2011;25(1):48–60.

21. Altizer S, Hobson KA, Davis AK, Roode JCD, Wassenaar LI. Do Healthy Monarchs Migrate Farther? Tracking Natal Origins of Parasitized vs. Uninfected Monarch Butterflies Overwintering in Mexico. PLOS ONE. 2015 Nov 25;10(11):e0141371.

22. Peacock SJ, Bouhours J, Lewis MA, Molnár PK. Macroparasite dynamics of migratory host populations. Theor Popul Biol. 2018 Mar 1;120:29–41.

23. Ghai RR, Fugère V, Chapman CA, Goldberg TL, Davies TJ. Sickness behaviour associated with non-lethal infections in wild primates. Proc R Soc B Biol Sci. 2015 Sep 7;282(1814):20151436.

24. Croft DP, Edenbrow M, Darden SK, Ramnarine IW, van Oosterhout C, Cable J. Effect of gyrodactylid ectoparasites on host behaviour and social network structure in guppies *Poecilia reticulata*. Behav Ecol Sociobiol. 2011 Dec 1;65(12):2219–27.

25. Kavaliers M, Colwell DD, Cloutier CJ, Ossenkopp KP, Choleris E. Pathogen threat and unfamiliar males rapidly bias the social responses of female mice. Anim Behav. 2014 Nov 1;97:105–11.

26. Ostfeld RS, Keesing F. Pulsed resources and community dynamics of consumers in terrestrial ecosystems. Trends Ecol Evol. 2000 Jun 1;15(6):232–7.

27. Touzot L, Schermer É, Venner S, Delzon S, Rousset C, Baubet É, et al. How does increasing mast seeding frequency affect population dynamics of seed consumers? Wild boar as a case study. Ecol Appl. 2020;30(6):e02134.

28. Prévot-Julliard AC, Henttonen H, Yoccoz NG, Stenseth NChR. Delayed maturation in female bank voles: optimal decision or social constraint? J Anim Ecol. 1999;68(4):684–97.

29. Bondrup-Nielsen S, Karlsson F. Movements and spatial patterns in populations of *Clethrionomys* species: A review. Ann Zool Fenn. 1985;22(3):385–92.

30. Grear DA, Perkins SE, Hudson PJ. Does elevated testosterone result in increased exposure and transmission of parasites? Ecol Lett. 2009;12(6):528–37.

31. Godfrey SS. Networks and the ecology of parasite transmission: A framework for wildlife parasitology. Int J Parasitol Parasites Wildl. 2013 Dec 1;2:235–45.

32. Craft ME, Caillaud D. Vol. 2011, Interdisciplinary Perspectives on Infectious Diseases. Hindawi; 2011 [cited 2020 May 15]. p. e676949 Network Models: An Underutilized Tool in Wildlife Epidemiology? Available from: https://www.hindawi.com/journals/ipid/2011/676949/

33. Keeling MJ, Eames KTD. Networks and epidemic models. J R Soc Interface. 2005 Sep 22;2(4):295–307.

34. Silk MJ, Croft DP, Delahay RJ, Hodgson DJ, Boots M, Weber N, et al. Using Social Network Measures in Wildlife Disease Ecology, Epidemiology, and Management. BioScience. 2017 Mar 1;67(3):245–57.

35. Corner LAL, Pfeiffer DU, Morris RS. Social-network analysis of *Mycobacterium bovis* transmission among captive brushtail possums (Trichosurus vulpecula). Prev Vet Med. 2003 Jun 12;59(3):147–67.

36. VanderWaal KL, Atwill ER, Hooper S, Buckle K, McCowan B. Network structure and prevalence of *Cryptosporidium* in Belding’s ground squirrels. Behav Ecol Sociobiol. 2013 Dec 1;67(12):1951–9.

37. Drewe JA. Who infects whom? Social networks and tuberculosis transmission in wild meerkats. Proc R Soc B Biol Sci. 2010 Feb 22;277(1681):633–42.

38. Powell SN, Wallen MM, Miketa ML, Krzyszczyk E, Foroughirad V, Bansal S, et al. Sociality and tattoo skin disease among bottlenose dolphins in Shark Bay, Australia. Behav Ecol. 2020 Mar 20;31(2):459–66.

39. VanderWaal KL, Atwill ER, Isbell LA, McCowan B. Linking social and pathogen transmission networks using microbial genetics in giraffe (*Giraffa camelopardalis*). J Anim Ecol. 2014;83(2):406–14.

40. Perkins SE, Ferrari MF, Hudson PJ. The effects of social structure and sex-biased transmission on macroparasite infection. Parasitology. 2008 Nov;135(13):1561–9.

41. White LA, Forester JD, Craft ME. Using contact networks to explore mechanisms of parasite transmission in wildlife. Biol Rev. 2017;92(1):389–409.

42. Bull CM, Godfrey SS, Gordon DM. Social networks and the spread of *Salmonella* in a sleepy lizard population. Mol Ecol. 2012;21(17):4386–92.

43. Fenner AL, Godfrey SS, Bull CM. Using social networks to deduce whether residents or dispersers spread parasites in a lizard population. J Anim Ecol. 2011;80(4):835–43.

44. Godfrey SS, Bull CM, James R, Murray K. Network structure and parasite transmission in a group living lizard, the gidgee skink, Egernia stokesii. Behav Ecol Sociobiol. 2009 May 1;63(7):1045–56.

45. Godfrey SS, Moore JA, Nelson NJ, Bull CM. Social network structure and parasite infection patterns in a territorial reptile, the tuatara (*Sphenodon punctatus*). Int J Parasitol. 2010 Nov 1;40(13):1575–85.

46. Leu ST, Kappeler PM, Bull CM. Refuge sharing network predicts ectoparasite load in a lizard. Behav Ecol Sociobiol. 2010 Sep 1;64(9):1495–503.

47. Craft ME. Infectious disease transmission and contact networks in wildlife and livestock. Philos Trans R Soc B Biol Sci. 2015 May 26;370(1669):20140107.

48. Forbes KM, Mappes T, Sironen T, Strandin T, Stuart P, Meri S, et al. Food limitation constrains host immune responses to nematode infections. Biol Lett. 2016 Sep 30;12(9):20160471.

49. Sweeny AR, Clerc M, Pontifes PA, Venkatesan S, Babayan SA, Pedersen AB. Supplemented nutrition decreases helminth burden and increases drug efficacy in a natural host–helminth system. Proc R Soc B Biol Sci. 2021 Jan 27;288(1943):20202722.

50. Haukisalmi V, Henttonen H. Variability of helminth assemblages and populations in the bank vole *Clethrionomys glareolus*. Pol J Ecol. 2000 Jan 1;48:219–31.

51. Khalil H, Ecke F, Evander M, Bucht G, Hörnfeldt B. Population Dynamics of Bank Voles Predicts Human Puumala Hantavirus Risk. EcoHealth. 2019 Sep 1;16(3):545–57.

52. Voutilainen L, Kallio ER, Niemimaa J, Vapalahti O, Henttonen H. Temporal dynamics of Puumala hantavirus infection in cyclic populations of bank voles. Sci Rep. 2016 Feb 18;6(1):1–15.

53. Forbes KM, Sironen T, Plyusnin A. Hantavirus maintenance and transmission in reservoir host populations. Curr Opin Virol. 2018 Feb 1;28:1–6.

54. Kallio ER, Klingström J, Gustafsson E, Manni T, Vaheri A, Henttonen H, et al. Prolonged survival of Puumala hantavirus outside the host: evidence for indirect transmission via the environment. J Gen Virol. 2006;87(8):2127–34.

55. Mazurkiewicz M. Shape, size and distribution of home ranges of Clethrionomys glareolus (Schreber, 1780). Acta Theriol (Warsz). 1971;16(2):23–60.

56. Mazurkiewicz M. Spatial organization of the population. Acta Theriol (Warsz). 1983;28(Suppl. 1):103–44.

57. Tamarin RH, Ostfeld RS, Pugh SR, Bujalska G. Social Systems and Population Cycles in Voles. Birkhäuser Verlag; 1990. 237 p.

58. Mistrick J, Veitch JSM, Kitchen SM, Clague S, Newman BC, Hall RJ, et al. Effects of food supplementation and helminth removal on space use and spatial overlap in wild rodent populations. J Anim Ecol. 2024;93(6):743–54.

59. Anonymous. Details omitted for double anonymised reviewing. 2024.

60. Clerc M, Babayan SA, Fenton A, Pedersen AB. Age affects antibody levels and anthelmintic treatment efficacy in a wild rodent. Int J Parasitol Parasites Wildl. 2019 Apr 1;8:240–7.

61. Kallio-Kokko H, Laakkonen J, Rizzoli A, Tagliapietra V, Cattadori I, Perkins SE, et al. Hantavirus and arenavirus antibody prevalence in rodents and humans in Trentino, Northern Italy. Epidemiol Infect. 2006 Aug;134(4):830–6.

62. Meyer BJ, Schmaljohn CS. Persistent hantavirus infections: characteristics and mechanisms. Trends Microbiol. 2000 Feb 1;8(2):61–7.

63. Kallio ER, Poikonen A, Vaheri A, Vapalahti O, Henttonen H, Koskela E, et al. Maternal antibodies postpone hantavirus infection and enhance individual breeding success. Proc R Soc B Biol Sci. 2006 Nov 7;273(1602):2771–6.

64. Pritchard MH, Kruse G. The collection and preservation of animal parasites. Tech. Bull. no. 1 [Internet]. Lincoln, Nebraska: Harold W. Manter Laboratory.: Lincoln, Nebraska: Harold W. Manter Laboratory. University of Nebraska Press; 1982 [cited 2023 Nov 16]. Available from: https://cir.nii.ac.jp/crid/1130000795640263296

65. Bates D, Maechler M, Bolker B, Walker S, Christensen R, Singmann H, et al. Package “lme4.” Convergence. 2015;12(1):2.

66. Kuznetsova A, Brockhoff PB, Christensen RHB. lmerTest Package: Tests in Linear Mixed Effects Models. J Stat Softw. 2017 Dec 6;82:1–26.

67. Kaikusalo A. Population turnover and wintering of the bank vole, *Clethrionomys glareolus* (Schreb.), in southern and central Finland. Ann Zool Fenn. 1972;9(4):219–24.

68. Haukisalmi V, Henttonen H, Tenora F. Population Dynamics of Common and Rare Helminths in Cyclic Vole Populations. J Anim Ecol. 1988;57(3):807–25.

69. Myllymäki A. Demographic Mechanisms in the Fluctuating Populations of the Field Vole *Microtus agrestis*. Oikos. 1977;29(3):468–93.

70. Myllymäki A. Intraspecific Competition and Home Range Dynamics in the Field Vole *Microtus agrestis*. Oikos. 1977;29(3):553–69.

71. Henttonen H. Importance of demography in understanding disease ecology in small mammals. THERYA. 2022 Jan 4;13(1):33.

72. Wanelik KM, Farine DR. A new method for characterising shared space use networks using animal trapping data. Behav Ecol Sociobiol. 2022 Aug 26;76(9):127.

73. Csardi G, Nepusz T. The Igraph Software Package for Complex Network Research. InterJournal Complex Syst. 2006;1695(5):1–9.

74. Farine DR, Whitehead H. Constructing, conducting and interpreting animal social network analysis. J Anim Ecol. 2015;84(5):1144–63.

75. Strandin T, Smura T, Ahola P, Aaltonen K, Sironen T, Hepojoki J, et al. Orthohantavirus Isolated in Reservoir Host Cells Displays Minimal Genetic Changes and Retains Wild-Type Infection Properties. Viruses. 2020 Apr;12(4):457.

76. R Core Team. R: A language and environment for statistical computing. [Internet]. Vienna, Austria: R Foundation for Statistical Computing. Vienna, Austria.; 2023. Available from: URL https://www.R-project.org/

77. Pessanha TS, Herrera HM, Jansen AM, Iñiguez AM. “Mi Casa, Tu Casa”: the coati nest as a hub of *Trypanosoma cruzi* transmission in the southern Pantanal biome revealed by molecular blood meal source identification in triatomines. Parasit Vectors. 2023 Jan 23;16(1):26.

78. Garnett BT, Delahay RJ, Roper TJ. Use of cattle farm resources by badgers (*Meles meles*) and risk of bovine tuberculosis (*Mycobacterium bovis*) transmission to cattle. Proc R Soc Lond B Biol Sci. 2002 Jul 22;269(1499):1487–91.

79. Titcomb G, Mantas JN, Hulke J, Rodriguez I, Branch D, Young H. Water sources aggregate parasites with increasing effects in more arid conditions. Nat Commun. 2021 Dec 3;12(1):7066.

80. Robert K, Garant D, Pelletier F. Keep in touch: Does spatial overlap correlate with contact rate frequency? J Wildl Manag. 2012;76(8):1670–5.

81. Becker DJ, Hall RJ. Too much of a good thing: resource provisioning alters infectious disease dynamics in wildlife. Biol Lett. 2014 Jul 31;10(7):20140309.

82. Ezenwa VO, Jolles AE. From Host Immunity to Pathogen Invasion: The Effects of Helminth Coinfection on the Dynamics of Microparasites. Integr Comp Biol. 2011 Oct 1;51(4):540–51.

83. Budischak SA, Sakamoto K, Megow LC, Cummings KR, Urban JF, Ezenwa VO. Resource limitation alters the consequences of co-infection for both hosts and parasites. Int J Parasitol. 2015 Jun 1;45(7):455–63.

84. Albery GF, Kirkpatrick L, Firth JA, Bansal S. Unifying spatial and social network analysis in disease ecology. J Anim Ecol. 2021;90(1):45–61.

85. Anonymous. Details omitted for double anonymous peer review. Dryad; 2024.

86. Bujalska G. Social System of the Bank Vole, Clethrionomys glareolus. In: Tamarin RH, Ostfeld RS, Pugh SR, Bujalska G, editors. Social Systems and Population Cycles in Voles [Internet]. Basel: Birkhäuser; 1990 [cited 2023 Jun 26]. p. 155–67. (Advances in Life Sciences). Available from: 10.1007/978-3-0348-6416-9_15

