## Supplemental materials for "Ecological factors alter how spatial overlap predicts viral infection dynamics in wild rodent populations"

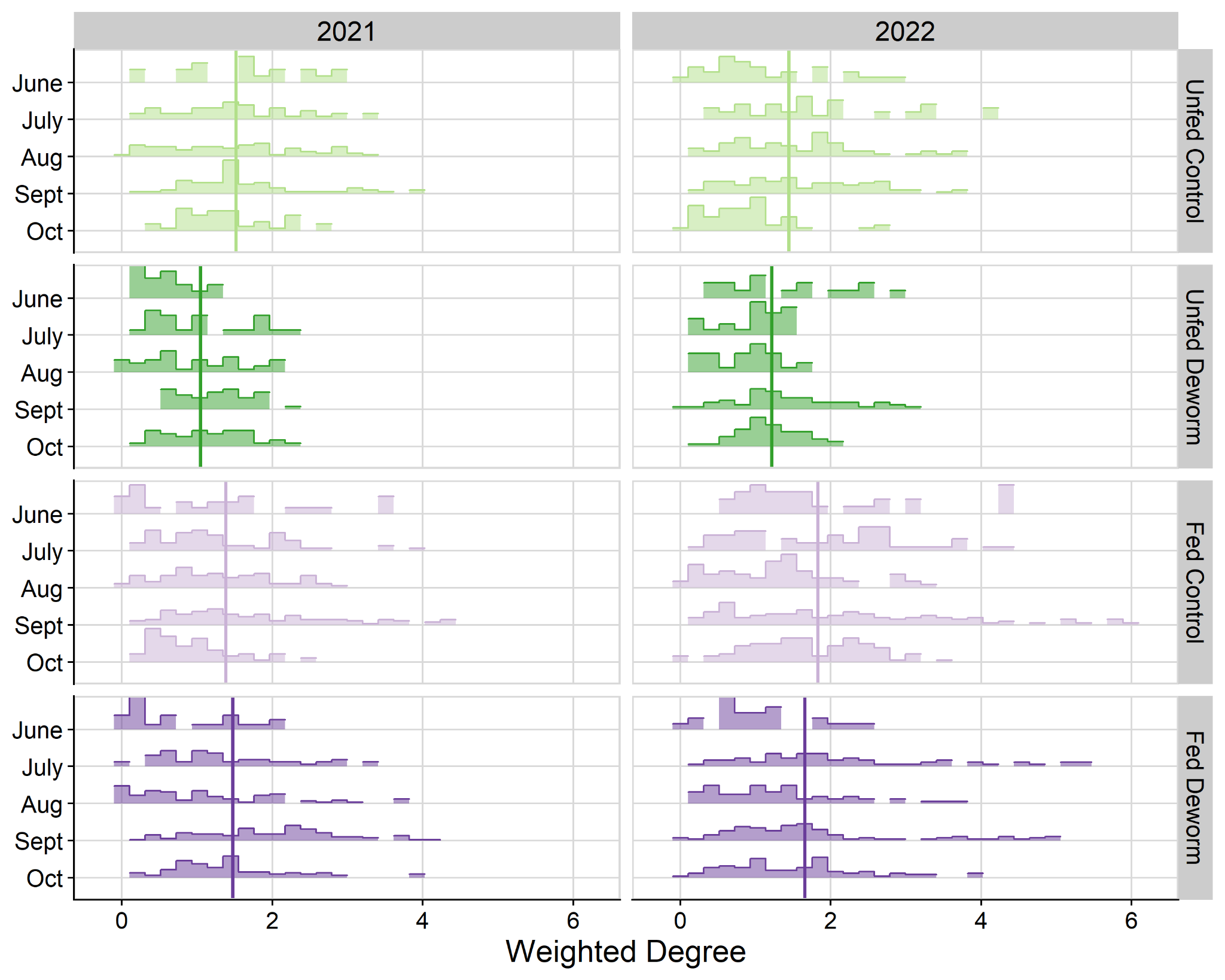

**Figure S1.** Weighted degree of bank voles in observed spatial overlap networks by month and treatment, faceted by study year. Weighted degree is the sum of all the pairwise space-use overlap between a focal vole and each of its neighbours - indicating a vole’s total spatial overlap with conspecifics in a given month. Vertical bars indicate the mean weighted degree per treatment, across all months in a given year.

**Exploratory behaviour**

*Methods*

In addition to spatial overlap, we quantified the tendency for voles to visit different traps as a measure of exploratory behaviour. Exploratory behaviour can influence environmental exposure and direct interactions in ways that may not be captured in spatial overlap. Exploratory behaviour was measured as the number of different traps a vole was caught in, relative to the expected number of traps for that number of captures (as described by 36). A power regression model $\left( y=0.98x^{0.74} \right)$ was fitted to the observed data of the number of unique traps and the number of captures per vole across a sampling year, one point for each vole in our dataset. A vole’s measure of exploratory behaviour was the residual of its observed number of unique traps minus the expected number of traps for a vole that was captured that many times, based on the model. Positive values indicate an individual used more unique traps than expected, whereas negative values indicate an animal used fewer unique traps than expected (36).

*Results*

Exploratory behaviour was negatively correlated with current infection in both the fed-control and fed-deworm treatments, indicating more exploratory animals were less likely to be infected (F-C: OR=0.58, CI [0.37-0.91], p=0.018; F-D: OR=0.63, CI [0.42-0.96], p=0.029). Exploratory behaviour was not correlated with current infection in either of the unfed treatments (p>0.40).

**Table S1.** Candidate models fitted by treatment. Boxes marked with an “X” indicate the models (columns) that were successfully fitted for each treatment (rows). Twelve potential candidate models were considered: Four degree measures: weighted degree, weighted (“Wt.”) degree partitioned by sex; weighted degree by reproductive status (“Repro”); weighted degree by functional group (“Fxnl grp”; combination of sex and reproductive status) were used as explanatory variables, each in separate models. For each degree measure, three models were fitted with interaction effects between weighted degree and each of sex, reproductive status, and both sex and reproductive status.

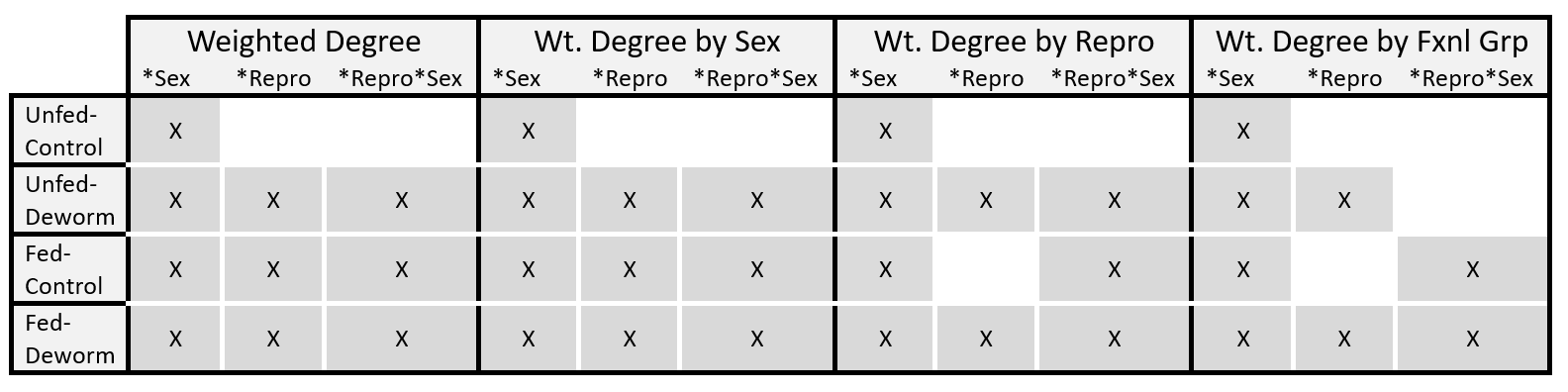

**Table S2.** Model summary for generalised linear mixed-effects model predicting current hantavirus infection status for voles in the unfed-control treatment. Odds ratios (OR) and 95% confidence intervals (CI) are reported for all predictors with reference levels listed first. Significant p-values are bolded. Site ID was included as a random effect.

| **Variable** | **OR** | **95% CI** | **p-value** |
| --- | --- | --- | --- |
| **Sex** |  |  |  |
| *Female* |  |  |  |
| *Male* | 0.40 | 0.04, 3.91 | 0.43 |
| **Exploratory Behaviour** | 1.26 | 0.74, 2.13 | 0.40 |
| **Previous Month** | 0.67 | 0.31, 1.47 | 0.32 |
| **Previous Network Size** | 1.50 | 0.32, 7.01 | 0.60 |
| **Year** |  |  |  |
| *2021* |  |  |  |
| *2022* | 1.07 | 0.39, 2.95 | 0.90 |
| **Repro Degree * Sex** |  |  |  |
| *Repro Degree * Female* | 0.86 | 0.29, 2.53 | 0.78 |
| *Repro Degree * Male* | 1.74 | 1.07, 2.82 | **0.025** |
| **Non-Repro Degree * Sex** |  |  |  |
| *Non-Repro Degree * Female* | 0.44 | 0.06, 3.52 | 0.44 |
| *Non-Repro Degree * Male* | 0.05 | 0.00, 1.41 | **0.079** |

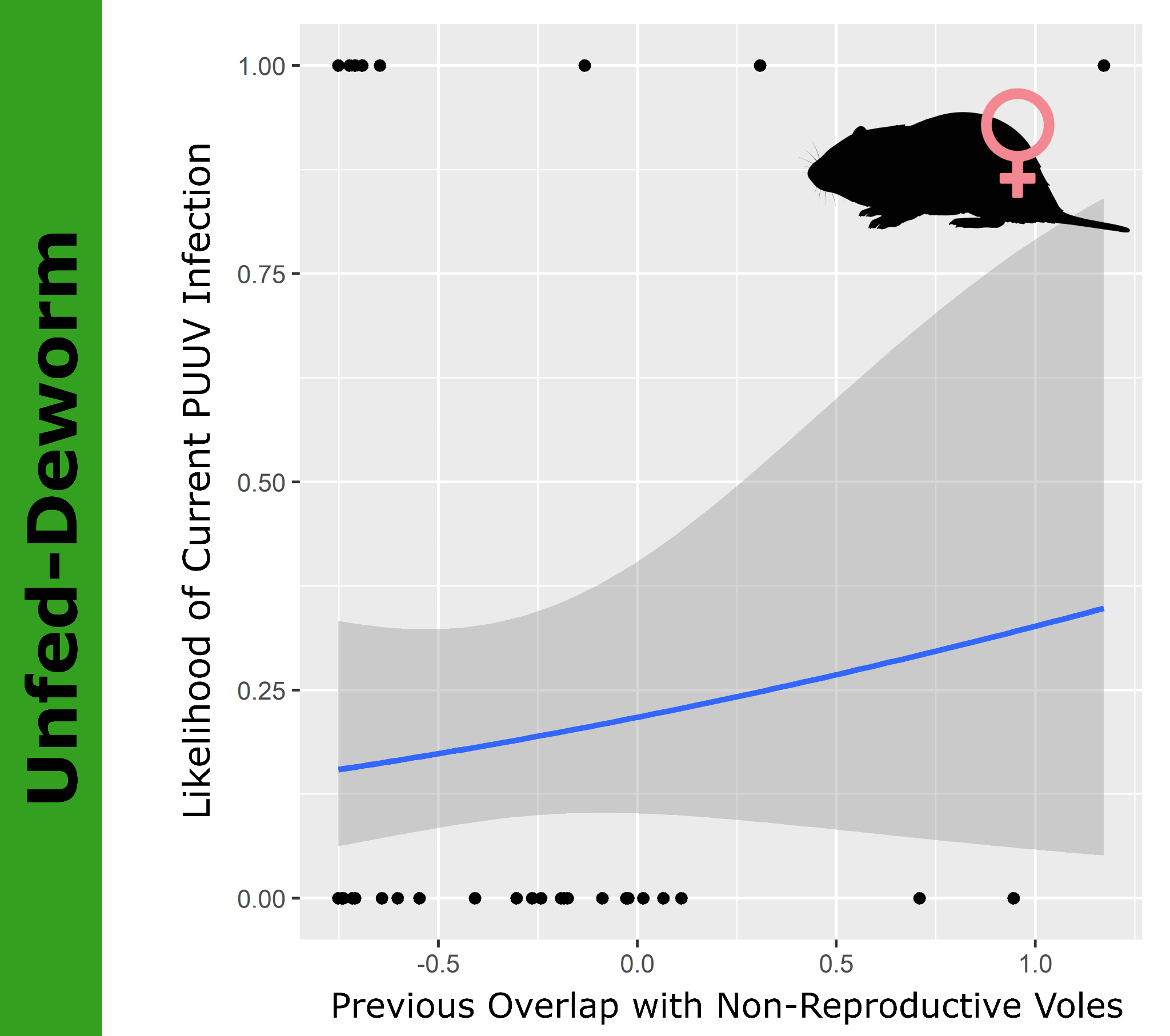

**Figure S2.** Correlation between likelihood of current Puumala hantavirus (PUUV) infection and previous spatial overlap (where overlap values were scaled and centred) for female voles in the unfed-deworm treatment. Previous overlap with non-reproductive voles increased infection likelihood. Points indicate raw data.

**Table S3.** Model summary for generalised linear mixed-effects model predicting current hantavirus infection status for voles in the unfed-deworm treatment. Odds ratios (OR) and 95% confidence intervals (CI) are reported for all predictors with reference levels listed first. Significant p-values are bolded. Site ID was included as a random effect.

| **Variable** | **OR** | **95% CI** | **p-value** |
| --- | --- | --- | --- |
| **Sex** |  |  |  |
| *Female* |  |  |  |
| *Male* | 1.03 | 0.16, 6.69 | 0.97 |
| **Reproductive Status (seasonal)** |  |  |  |
| *Reproductive* |  |  |  |
| *Non-Reproductive* | 0.68 | 0.11, 4.02 | 0.67 |
| **Exploratory Behaviour** | 1.04 | 0.53, 2.04 | 0.90 |
| **Previous Month** | 0.31 | 0.13, 0.78 | **0.012** |
| **Previous Network Size** | 0.42 | 0.02, 8.93 | 0.58 |
| **Year** |  |  |  |
| *2021* |  |  |  |
| *2022* | 0.30 | 0.08, 1.11 | 0.071 |
| **Repro Degree * Sex** |  |  |  |
| *Repro Degree * Female* | 0.84 | 0.12, 5.86 | 0.86 |
| *Repro Degree * Male* | 1.60 | 0.37, 6.79 | 0.53 |
| **Non-Repro Degree * Sex** |  |  |  |
| *Non-Repro Degree * Female* | 21.3 | 1.72, 266 | **0.017** |
| *Non-Repro Degree * Male* | 1.89 | 0.37, 9.51 | 0.44 |

**Table S4.** Model summary for generalised linear mixed-effects model predicting current hantavirus infection status for voles in the fed-control treatment. Odds ratios (OR) and 95% confidence intervals (CI) are reported for all predictors with reference levels listed first. Significant p-values are bolded. Site ID was included as a random effect.

| **Variable** | **OR** | **95% CI** | **p-value** |
| --- | --- | --- | --- |
| **Sex** |  |  |  |
| *Female* |  |  |  |
| *Male* | 2.06 | 0.59, 7.14 | 0.26 |
| **Reproductive Status (seasonal)** |  |  |  |
| *Reproductive* |  |  |  |
| *Non-Reproductive* | 0.00 | 0.00, 0.19 | **0.009** |
| **Exploratory Behaviour** | 0.58 | 0.37, 0.91 | **0.018** |
| **Previous Month** |  |  |  |
| *June* |  |  |  |
| *July* | 1.31 | 0.20, 8.46 | 0.78 |
| *August* | 0.89 | 0.12, 6.46 | 0.91 |
| *September* | 0.18 | 0.02, 2.02 | 0.16 |
| **Previous Network Size** | 1.27 | 0.54, 2.99 | 0.59 |
| **Year** |  |  |  |
| *2021* |  |  |  |
| *2022* | 0.04 | 0.01, 0.15 | **<0.001** |
| **Repro Male Degree * ReproStatus * Sex** |  |  |  |
| *R M Degree * Repro * Female* | 0.72 | 0.25, 2.09 | 0.55 |
| *R M Degree * Repro * Male* | 1.65 | 0.86, 3.16 | 0.13 |
| *R M Degree * Non-Repro * Female* | 4.21 | 0.42, 42.6 | 0.22 |
| *R M Degree * Non-Repro * Male* | 0.03 | 0.00, 6.83 | 0.20 |
| **Non-Repro Male Degree * ReproStatus * Sex** | |  |  |
| *N-R M Degree * Repro * Female* | 2.81 | 1.13, 6.97 | **0.026** |
| *N-R M Degree * Repro * Male* | 0.93 | 0.27, 3.15 | 0.91 |
| *N-R M Degree * Non-Repro * Female* | 3.05 | 0.40, 23.1 | 0.28 |
| *N-R M Degree * Non-Repro * Male* | 4,549 | 0.58, 35,797,396 | 0.066 |
| **Repro Female Degree * ReproStatus * Sex** | |  |  |
| R F Degree * Repro * Female | 0.73 | 0.32, 1.66 | 0.45 |
| R F Degree * Repro * Male | 1.22 | 0.63, 2.36 | 0.55 |
| R F Degree * Non-Repro * Female | 0.00 | 0.00, 0.10 | **0.009** |
| R F Degree * Non-Repro * Male | 0.00 | 0.00, 0.25 | **0.018** |
| **Non-Repro Female Degree * ReproStatus * Sex** | |  |  |
| *N-R F Degree * Repro * Female* | 0.86 | 0.41, 1.79 | 0.68 |
| *N-R F Degree * Repro * Male* | 0.55 | 0.19, 1.59 | 0.27 |
| *N-R F Degree * Non-Repro * Female* | 0.37 | 0.08, 1.59 | 0.18 |
| *N-R F Degree * Non-Repro * Male* | 7.97 | 0.57, 111 | 0.12 |

**Fed-Control Treatment**

*Results*

In the fed-control treatment, non-reproductive male and female voles with previous high spatial overlap with female breeders were less likely to be currently infected (Males: OR=0.0, p=0.01; Females: OR=0.0, p=0.019; **Figure S2**). However, the sample size of infected non-reproductive voles was very low (males n=2; females n=4) limiting our ability to draw robust conclusions from these findings.

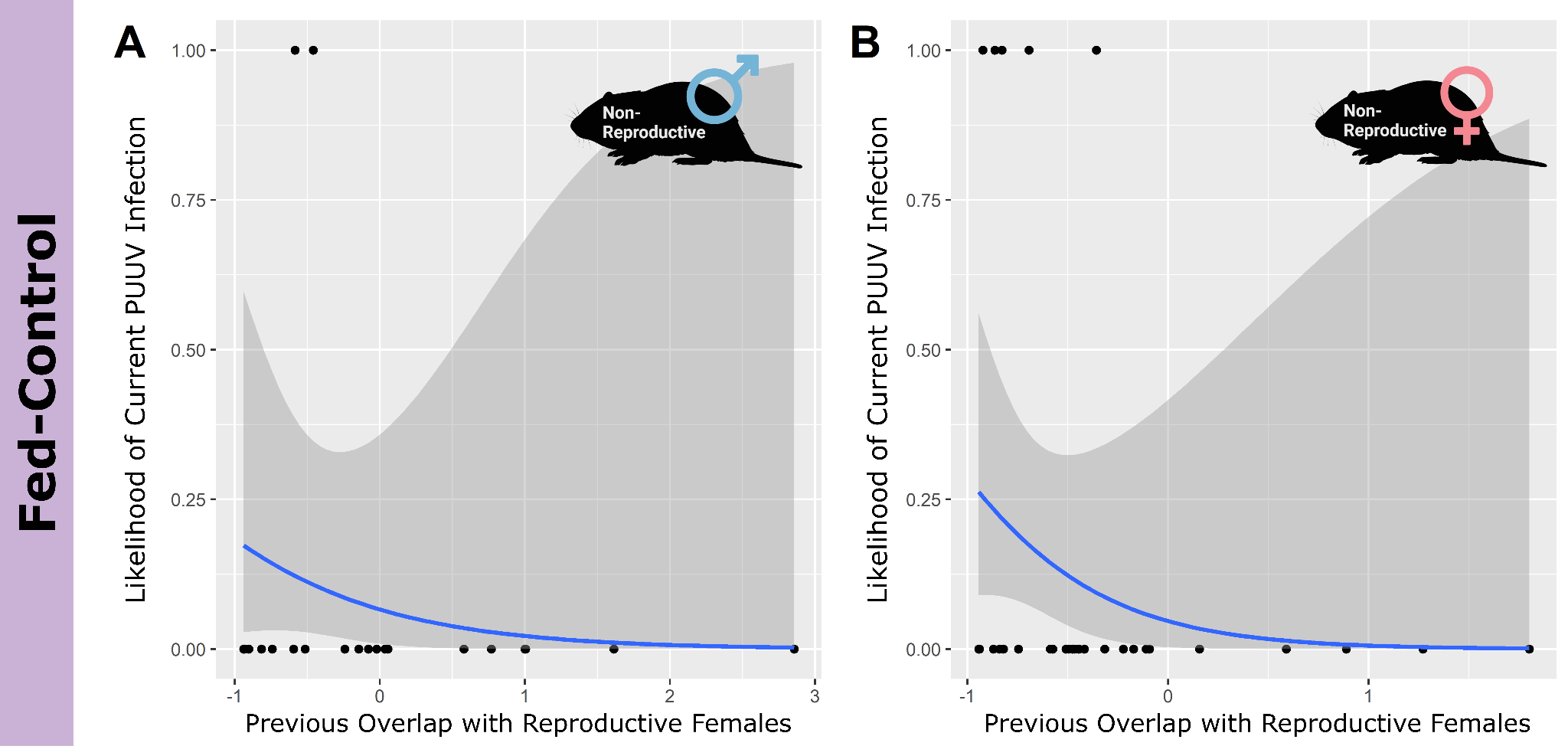

**Figure S3.** Correlation between likelihood of current Puumala hantavirus (PUUV) infection and previous spatial overlap (where overlap values were scaled and centred) in the fed-control treatment. For both A) non-reproductive male voles and B) non-reproductive female voles, previous overlap with reproductive female voles decreased the likelihood of current infection. Points indicate raw data.

*Discussion*

Young bank voles may remain in their mother’s territory before dispersing to establish their own territory (86). Our finding that increased overlap with reproductive females may decrease infection likelihood in non-reproductive individuals could indicate a protective effect of remaining in natal territory. Non-reproductive voles with high spatial overlap with reproductive females may not explore widely themselves, limiting their exposure to environmental pathogens.

**Table S5.** Model summary for generalised linear mixed-effects model predicting current hantavirus infection status for voles in the fed-deworm treatment. Odds ratios (OR) and 95% confidence intervals (CI) are reported for all predictors with reference levels listed first. Significant p-values are bolded. Site ID was included as a random effect.

| **Characteristic** | **OR** | **95% CI** | **p-value** |
| --- | --- | --- | --- |
| **Sex** |  |  |  |
| *Female* |  |  |  |
| *Male* | 4.01 | 1.49, 10.8 | **0.006** |
| **Reproductive Status (seasonal)** |  |  |  |
| *Reproductive* |  |  |  |
| *Non-Reproductive* | 0.01 | 0.00, 0.75 | **0.037** |
| **Exploratory Behaviour** | 0.63 | 0.42, 0.96 | **0.029** |
| **Previous Month** | 1.58 | 0.78, 3.18 | 0.20 |
| **Previous Network Size** | 0.44 | 0.17, 1.14 | 0.090 |
| **Year** |  |  |  |
| *2021* |  |  |  |
| *2022* | 3.20 | 1.03, 9.89 | **0.043** |
| **Repro Male Degree * ReproStatus * Sex** |  |  |  |
| *R M Degree * Repro * Female* | 0.29 | 0.11, 0.79 | **0.015** |
| *R M Degree * Repro * Male* | 1.63 | 0.89, 3.00 | 0.11 |
| *R M Degree * Non-Repro * Female* | 4.84 | 0.08, 296 | 0.45 |
| *R M Degree * Non-Repro * Male* | 0.03 | 0.00, 3.94 | 0.16 |
| **Non-Repro Male Degree * ReproStatus * Sex** | | |  |
| *N-R M Degree * Repro * Female* | 1.81 | 0.79, 4.16 | 0.16 |
| *N-R M Degree * Repro * Male* | 0.76 | 0.40, 1.44 | 0.40 |
| *N-R M Degree * Non-Repro * Female* | 0.05 | 0.00, 37.4 | 0.38 |
| *N-R M Degree * Non-Repro * Male* | 1.64 | 0.73, 3.65 | 0.23 |
| **Repro Female Degree * ReproStatus * Sex** | | |  |
| R F Degree * Repro * Female | 4.22 | 1.54, 11.6 | **0.005** |
| R F Degree * Repro * Male | 0.78 | 0.45, 1.34 | 0.36 |
| R F Degree * Non-Repro * Female | 5.99 | 0.03, 1,236 | 0.51 |
| R F Degree * Non-Repro * Male | 1.54 | 0.36, 6.56 | 0.56 |
| **Non-Repro Female Degree * ReproStatus * Sex** | | |  |
| *N-R F Degree * Repro * Female* | 0.97 | 0.28, 3.39 | 0.96 |
| *N-R F Degree * Repro * Male* | 0.71 | 0.31, 1.62 | 0.42 |
| *N-R F Degree * Non-Repro * Female* | 0.00 | 0.00, 79.9 | 0.20 |
| *N-R F Degree * Non-Repro * Male* | 0.87 | 0.15, 5.18 | 0.88 |
